# Closing the gap in the clinical adoption of computational pathology: a standardized, open-source framework to integrate deep-learning algorithms into the laboratory information system

**DOI:** 10.1101/2024.07.11.603091

**Authors:** Miriam Angeloni, Davide Rizzi, Simon Schoen, Alessandro Caputo, Francesco Merolla, Arndt Hartmann, Fulvia Ferrazzi, Filippo Fraggetta

## Abstract

Digital pathology (DP) has revolutionized cancer diagnostics, allowing the development of deep-learning (DL) models supporting pathologists in their daily work and contributing to the improvement of patient care. However, the clinical adoption of such models remains challenging. Here we describe a proof-of-concept framework that, leveraging open-source DP software and Health Level 7 (HL7) standards, allows the integration of DL models in the clinical workflow.

Development and testing of the workflow were carried out in a fully digitized Italian pathology department. A Python-based server-client architecture was implemented to interconnect the anatomic pathology laboratory information system (AP-LIS) with an external artificial intelligence decision support system (AI-DSS) containing 16 pre-trained DL models through HL7 messaging. Open-source toolboxes for DL model deployment, including WSInfer and WSInfer-MIL, were used to run DL model inference. Visualization of model predictions as colored heatmaps was performed in QuPath.

As soon as a new slide is scanned, DL model inference is automatically run on the basis of the slide’s tissue type and staining. In addition, pathologists can initiate the analysis on-demand by selecting a specific DL model from the virtual slides tray. In both cases the AP-LIS transmits an HL7 message to the AI-DSS, which processes the message, runs DL model inference, and creates the appropriate type of colored heatmap on the basis of the employed classification model. The AI-DSS transmits model inference results to the AP-LIS, where pathologists can visualize the output in QuPath and/or directly from the virtual slides tray. The developed framework supports multiple DL toolboxes and it is thus suitable for a broad range of applications. In addition, this integration workflow is a key step to enable the future widespread adoption of DL models in pathology diagnostics.

## Introduction

The advent of digital pathology (DP), with the digitization of histopathology glass specimens into high-resolution whole-slide images (WSIs), revolutionized cancer diagnostics and research.^1^ The advantages offered by WSIs are manifold, including an improved laboratory workflow through easier archiving and retrieval of the virtual slides, as well as a seamless integration of patient data within the laboratory information system (LIS). Virtual slides additionally allow for remote consultation, which makes them ideal for rapid second opinions, teaching, and training.^2^ Importantly, the digitization of glass slides, coupled with the exponential improvements over the past decades in both hardware and software components, enabled the development and application of numerous artificial intelligence (AI)-based tools. This initiated a paradigm shift towards the adoption of an automated, accurate, computer-assisted diagnostic system.^3^ In precision oncology, AI tools have proven capable of tumor detection, subtyping, and classification, as well as more advanced applications such as prediction of genetic alterations and cancer prognosis.^4^

Over the past 10 years, the number of publications involving AI-based computational pathology (CPath) tools has increased more than 100-fold.^5^ Nevertheless, the vast majority of AI tools developed for the pathology domain, both commercial and non-commercial, are rarely adopted in routine diagnostics.^6^ There are several reasons for this. Firstly, the clinical adoption of AI tools requires a fully digital workflow. However, due to cultural and technical challenges,^7,8^ few anatomic pathology departments are fully digital. The lack of prospective clinical validation as well as the regulatory approval required for the clinical use of an AI-based diagnostic assay further limit AI implementation in the routine pathology practice^5,7^. In addition, the use of AI in routine diagnostics represents a financial burden for healthcare institutions as it is typically not yet reimbursed.^5,9^ Another major hurdle is the lack of an established protocol for integrating CPath solutions within existing anatomic pathology laboratory information systems (AP-LIS).^5,10^ Many commercial DL algorithms run on cloud servers, and often rely on proprietary viewers, whose integration within existing AP-LIS is non-trivial. Most of the publicly available DL models on the other hand are not reusable,^11,12^ not even in fully-digitized diagnostic workflows, nor accessible to pathologists without programming skills.^13^ Even for the minority of DL algorithms that have been published in a format suitable for use by other researchers, their integration into the AP-LIS remains challenging. Furthermore, the output of DL models often consists of class prediction probabilities that do not help pathologists understand the black-box nature of the AI solutions they are adopting. Thus, the integration of DL models into the diagnostic workflow represents a significant challenge yet to be addressed.^8,9^

The Caltagirone pathology department shifted towards a fully digitized diagnostic workflow in 2019, through the adoption of a lean approach and the implementation of a fully tracked pathology system integrated within the AP-LIS.^14,15^ Relying on the internationally recognized Health Level 7 (HL7) standard^16^, and the unique infrastructure offered by the Caltagirone pathology department, we developed the first standardized, open-source prototype workflow for integrating DL models into the AP-LIS. In this work, HL7 was used to interconnect the AI tools to the AP-LIS, and open-source DP software were employed both for inference of pre-trained models on routine WSIs and visualization of DL model predictions. Our results show how three key challenges limiting the adoption of CPath solutions into existing clinical workflows can be successfully addressed, namely (i) the establishment of an integration workflow, (ii) the inclusion of publicly available DL models, and (iii) the implementation of a visualization strategy for model inference results. This work is, to the best of our knowledge, the first proof-of-concept study aiming at bridging the gap in the clinical adoption of diagnostic and prognostic DL models, thereby marking an important step to move beyond academic research.

## Methods

### Caltagirone’s pathology department infrastructure

Development and testing of the integration workflow was carried out in a fully digitized Italian pathology laboratory at Gravina Hospital in Caltagirone, Italy.^15^ Here, all routine glass slides are scanned into whole-slide images (WSIs) with either a Panoramic P1000 scanner (3DHistech, Budapest, Hungary) or a Panoramic P250 scanner (3DHistech, Budapest, Hungary) at a resolution of 0.24 microns per pixel (mpp) and stored in the scanner manufacturer’s proprietary image format (.mrxs) on a Network Attached Storage (NAS) system (Qnap NAS TVS-EC1280U-SAS-RP) via a 100Mbit/s network connection.

The digital workflow uses the laboratory management application software Pathox (v13.32.0, Tesi Elettronica e Sistemi Informativi S.P.A., Milan, Italy) to implement a LIS-centric approach^17,18^, which enables WSIs to be opened directly from the virtual tray of the AP-LIS and visualized by default in the scanner-specific viewing software. In April 2023, the open-source bio-image analysis tool QuPath^19^ software was additionally integrated within the AP-LIS.

### Computational hardware and software

All computational tasks, including the implementation of the Python-based server-client architecture and deep-learning (DL) models deployment, were performed on a remote server located in the Gravina Hospital, equipped with two AMD EPYC 7313 processors and two AMD Radeon Instinct MI210 (64GB RAM) graphics processing units (GPUs) and relying on Ubuntu’s 22.04.4 long-term support (LTS) operating system.

The entire integration workflow was developed in a dedicated conda environment with Python v3.10.14. To leverage AMD GPUs, the PyTorch DL framework was installed with Radeon Open Compute (ROCm) support (v5.7) together with the libraries torch v2.3.1, and torchvision v0.18.1. For deployment of pre-trained DL models, one of the two GPUs was used. Pathologists’ workstations to visualize the output of DL model inference results included computers with consumer-grade monitors, a QuPath installation and hardware specifications as previously described.^15^

### Deployment of pre-trained deep-learning algorithms and results visualization in QuPath

Different types of DL models were integrated, including both strongly-supervised learning frameworks (hereafter referred to as ‘patch-level’ classification models) and weakly-supervised learning frameworks^20^ such as attention-based multiple-instance learning (MIL) approaches (hereafter referred to as ‘slide-level’ classification models).

Deployment of pre-trained patch-level classification models was performed relying on the WSInfer^13^ command-line (CLI) tool v0.6.1, installed in the dedicated conda environment via pip after PyTorch installation. Pre-trained weights for built-in models, with the associated configuration files, were retrieved relying on the Python package wsinfer-zoo v0.6.2 (https://github.com/SBU-BMI/wsinfer-zoo; https://zenodo.org/records/12680690), automatically installed as dependency during the WSInfer pip installation. Deployment of pre-trained slide-level classification models for the assessment of *TP53* mutational status and risk of cancer death was performed relying on the WSInfer-MIL CLI tool v0.1.0 (https://github.com/SBU-BMI/wsinfer-mil; https://zenodo.org/records/12680704). Deployment of models for the assessment of *BRAF* mutational status and microsatellite instability (MSI) was performed relying on the marugoto toolbox v0.8.0 (https://github.com/KatherLab/marugoto). The two pre-trained models were retrieved from the GitHub repository made public by Niehues et al.^21^ (https://github.com/KatherLab/crc-models-2022/tree/main/Quasar_models). For both classification tasks, the export-0.pkl file under the corresponding Wang+attMIL subfolder was chosen for inference.

Further details on the customization of the aforementioned toolboxes and the input parameters used for inference can be found in the Supplementary materials and methods.

Visualization of inference results from WSInfer and WSInfer-MIL CLI tools was performed by creating, for each analyzed WSI, a QuPath project storing a colored heatmap. To this aim, we relied on the open-source bio-image analysis software QuPath^19^ v0.4.3 and on the Python package paquo v0.8.0 (https://github.com/Bayer-Group/paquo copyright 2020 Bayer AG, licensed under GPL-3.0). Three different visualization styles were implemented, namely: measurement maps for binary patch-level classification tasks, color maps for multi-class patch-level classification tasks, and density maps for slide-level classification tasks. Detailed information on how each visualization style was implemented in paquo can be found in the Supplementary materials and methods.

### Interfacing between the AP-LIS and the AI-DSS through HL7 messaging

Hereafter we will refer to the set of DL models interconnected to the AP-LIS as external AI-based decision support system (AI-DSS). Interfacing between the AP-LIS and the AI-DSS relied on the internationally recognized Health Level 7 (HL7)^16^ version 2 protocol (v. 2.6) messaging. The communication flow between the AP-LIS and the AI-DSS, as well as the adopted messaging, followed the HL7 standard Pathox_rev1_5 specifications. In particular, all WSI analysis requests were transmitted from the AP-LIS to the AI-DSS via HL7 Laboratory Order Messages (OML^O33). Conversely, the inference results generated by the AI-DSS were transmitted to the AP-LIS via HL7 Unsolicited Laboratory Observation Messages (OUL^R21). In the implemented workflow, each OML^O33 message corresponds to the analysis request of a single WSI, even when multiple WSIs are available for the same patient. This ensures a single ‘specimen’ (SPM) segment and a single ‘observation request’/‘common order’ (OBR/ORC) segment pair. Information about the DL model to be used and the location of the WSI to analyze are stored respectively in the fields SPM.4.1^SPM.4.2 and OBR.13 of the OML^O33 message (**Supplementary Figure 1**). All incoming OML^O33 messages are placed in a queue and, before any further processing, an acknowledgment (ACK) message is transmitted to the AP-LIS. After the ACK message, processing of the analysis requests starts following the First-In-First-Out (FIFO) principle. For each message of the queue, first DL model’s name and WSI file location are retrieved, then inference is run with one of the installed CLI tools, and ultimately a QuPath project is generated to display DL model inference results as colored heatmap (**Figure 1**).

**Figure 1.**
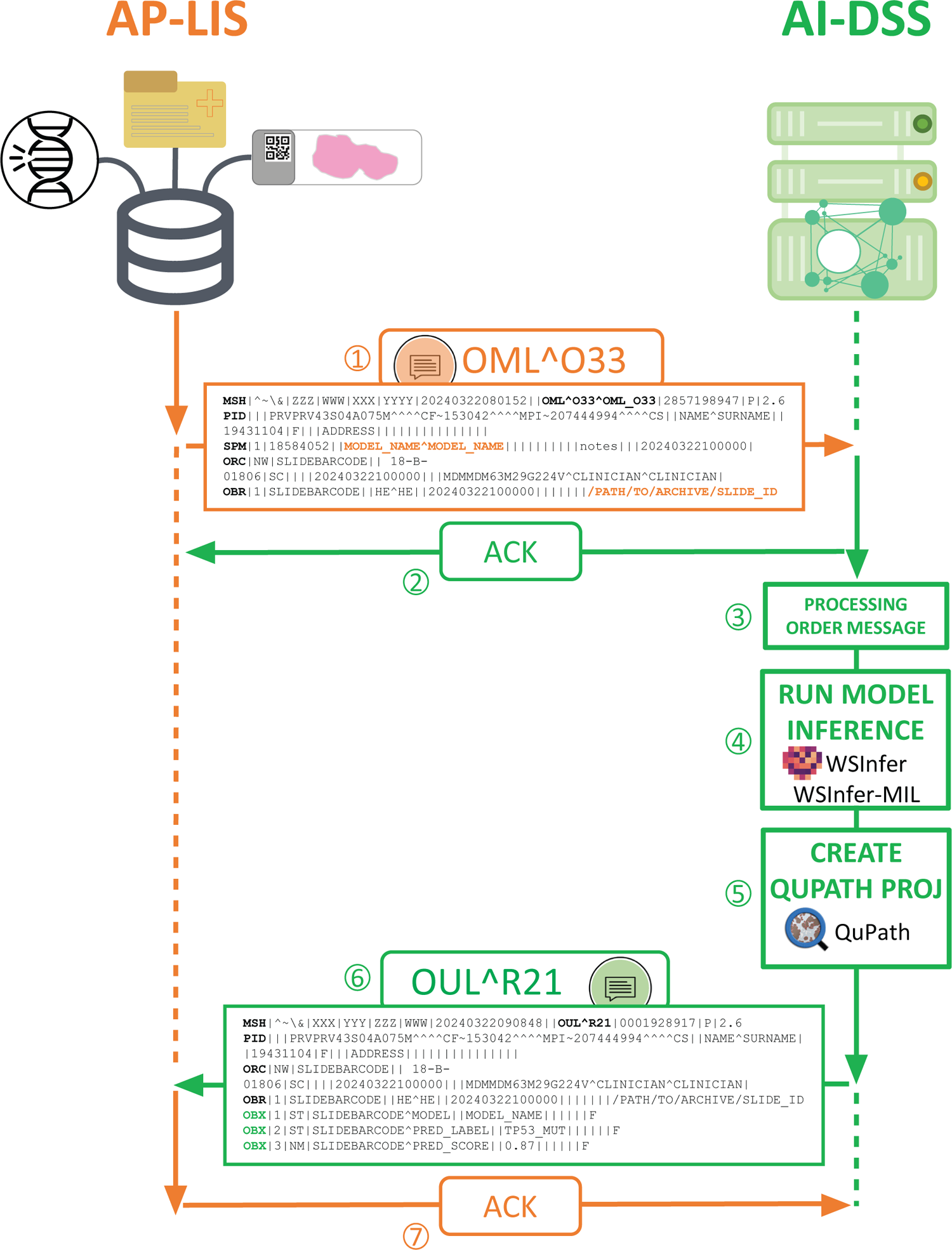
Schematization of the integration workflow. The anatomic pathology laboratory information system (AP-LIS) and the external artificial intelligence decision support system (AI-DSS) containing a set of pre-trained deep-learning (DL) models communicate with each other via Health Level 7 (HL7) messaging through the following sequence of events: (1) a laboratory order message (OML^O33) is sent from the AP-LIS to the AI-DSS; (2) an acknowledgment (ACK) message is sent from the AI-DSS to the AP-LIS upon reception of the OML^O33 order message; (3) the AI-DSS processes the OML^O33 order message retrieving information on the DL model name and the path to the WSI to analyze (bold orange); (4) DL model inference is run; (5) a QuPath project is built to visualize DL models results; (6) a laboratory observation message (OUL^R21) is sent from the AI-DSS to the AP-LIS to communicate DL model results as OBX segments (bold green); (7) an ACK message is sent from the AP-LIS to the AI-DSS upon reception of the OUL^R21 message.

Basing on the type of pre-trained DL model used for deployment, the number of ‘observation/result’ (OBX) segments included in the output OUL^R21 message may vary. The first OBX segment always contains a string indicating the name of the model. In addition, for all the WSIs where inference was performed with the WSInfer CLI tool, tissue mask (jpg file), tile-level predictions (CSV file), and metadata about the run (json file) are added as distinct OBX segments encoded in Base64 (**Supplementary Figure 2A**). For a selection of models, the top five tiles supporting the worst clinical outcome are Base64 encoded and added to the output OUL^R21 message as additional OBX segments (**Supplementary Figure 2B**). All slide-level classification models have instead, in addition to the first OBX segment, two OBX segments containing the slide-level predicted label (or risk class) and the associated prediction score (or risk score) (**Supplementary Figure 2C**).

After the transmission of the OUL^R21 HL7 message to the AP-LIS, the latter sends an ACK message to the AI-DSS following which the connection between the two is closed and the next message in the queue, if available, is processed. In the event of errors or failures, a maximum of three analysis attempts were programmed for the same slide. If none of the three attempts is successful, the next WSI in the queue, if available, is processed.

Communication between the AP-LIS and the AI-DSS was set-up via intranet connection using socket programming in a Python-based server-client architecture. The server-client architecture was developed to enable both server and client functionalities depending on the role of the system at any given time point. The server listens for incoming OML^O33 and ACK messages from the AP-LIS and, upon receiving a connection, reads and processes the incoming data. The client, on the other hand, transmits DL inference results as OUL^R21 messages and sends ACK messages to the AP-LIS. HL7 messaging exchange between the AP-LIS and the AI-DSS takes place via a Transmission Control Protocol/Internet Protocol (TCP/IP) connection using the Minimal Lower Layer Protocol (MLLP). The Python package Hl7apy v1.3.5 was used to analyze and generate HL7 messages.

## Results

Our integration workflow to incorporate both publicly available and custom developed deep-learning (DL) algorithms into the anatomic-pathology laboratory information system (AP-LIS) was tested in the fully digitized pathology department of Caltagirone, Italy.^15^ The workflow was designed around two core elements interconnected with each other: the AP-LIS and an external system that provides diagnostic support (**Figure 1**). Hereafter we will refer to this external system, which encompasses the set of DL models integrated within the AP-LIS, as to an AI-based decision support system (AI-DSS). Interconnection between the AP-LIS and the AI-DSS was set-up relying on a Python-based server-client architecture, and communication between the two systems relied on the internationally recognized Health Level 7 (HL7) messaging. Free and open-source resources were used both for running DL model deployment on routine digitized glass slides and for visualizing DL model inference results. Inference of DL models providing a patch-based classification was performed relying on WSInfer (<u>W</u>hole <u>S</u>lide <u>Infer</u>ence),^13^ which allows a streamlined deployment of both built-in and custom developed DL models in a one-line command. Instead, inference of DL models providing a slide-level classification, such as pre-trained multiple-instance learning (MIL) models, relied on WSInfer-MIL (https://github.com/SBU-BMI/wsinfer-mil; https://zenodo.org/records/12680704) and marugoto (https://github.com/KatherLab/marugoto). To allow pathologists a straightforward investigation and interpretation of DL model results, QuPath^19^ and its pythonic interface paquo (https://github.com/Bayer-Group/paquo) were employed to create, for each analyzed WSI, a QuPath project with DL model inference results displayed as colored heatmap overlaid to the original WSI.

Currently (last update 10.06.2024), a total of 16 algorithms can be applied to routine digitized hematoxylin and eosin (H&E) slides of the Caltagirone pathology department (**Table 1**). Eleven out of 16 DL models (N = 8 WSInfer built-in models; N = 3 in-house developed models) perform either a binary or a multi-class classification task to discriminate between different types of tissue (e.g., epithelium, stroma, tumor, necrosis, and dysplasia) or condition (e.g., tumor/non-tumor), while the remaining five are publicly available models to predict the status of predictive clinical biomarkers^21^ (i.e., *TP53*, *BRAF*, MSI) as well as the risk of cancer death in kidney renal papillary cell carcinoma (KIRP) and glioblastoma multiform and low-grade glioma (GMBLGG)^22^. Based on the performed classification task and whether or not an attention-based MIL approach was employed, three types of visualization styles were implemented in paquo and visualized as colored heatmap in QuPath: (i) measurement maps, (ii) color maps, and (iii) density maps (**Figure 2**). Irrespective of the visualization style, implementation was made so that, when opening the QuPath project associated with a given WSIs, a transparent tiled overlay is already superimposed on the original slide and the colored visualization can be turned on by pathologists manually from QuPath graphical interface. Measurement maps were used to visualize the output of binary classification tasks (e.g., tumor-positive/tumor-negative, metastasis/non-metastasis). In QuPath, measurement maps provide a color-coded representation of a measure associated with a given detection object. Here, as measure for each tile detection object we used the prediction score associated with the worst class label from the clinical point of view (e.g., tumor in tumor/non-tumor, metastasis in metastasis/non-metastasis) (**Figure 2A**). For measurement maps, pathologists can enable the colored visualization as shown in **Video 1**. Multi-class classification problems (e.g., classification into benign/grade3/grade4or5 in prostate tissue) were instead visualized through color maps. Here, the QuPath project was built to support a number of pre-defined annotation classes, each assigned a distinct color, matching the number of labels predicted by the multi-class DL model. Tiles within the tissue sample are colored according to the color of the annotation class corresponding to the predicted label (**Figure 2B**). For color maps, pathologists can enable the colored visualization as shown in **Video 2**. As visualization style for attention-based MIL models, density maps were implemented. Indeed, attention-based MIL algorithms provide in output an overall slide-level prediction score rather than a tile-level prediction score. On the other hand, attention scores are available at tile-level. Density maps in QuPath were originally conceived to help finding hotspots for specific clinical applications (e.g., Ki67 scoring), and more in general to find areas with high objects density. Here, we relied on the density heatmap to highlight tiles, and thus areas, associated with higher attention scores. As additional support to pathologists in interpreting the output of MIL algorithms, a summary of model’s predicted label and associated prediction score were provided as description of the QuPath project (**Figure 2C**). Density map visualization can be enabled in QuPath as shown in **Video3**. Taken together, the three implemented visualization styles allow pathologists to explore and get insights into DL model predictions, thus partially overcoming the black-box nature of the adopted AI-based algorithm.

**Figure 2.**
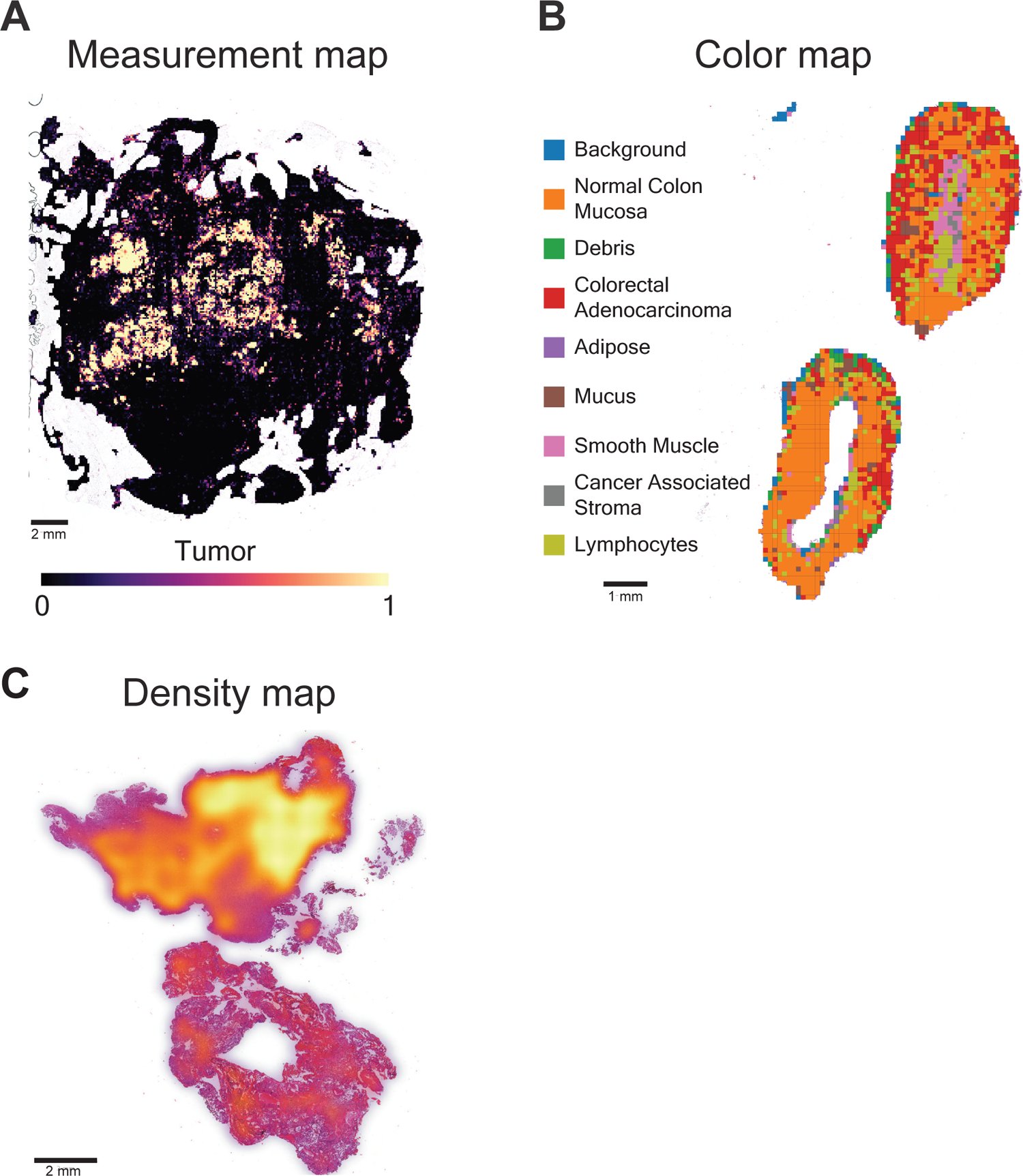
Visualization styles of deep-learning models output in QuPath. Exemplary visualization output of different classification models. (A) Measurement map for binary classification obtained by applying the ‘breast-tumor-resnet34.tcga-brca’ model on breast tissue. The measurement associated with a given tile refers to its prediction score for the class ‘Tumor’: the brighter the color, the higher the prediction score (basing on the chosen color scale). (B) Color map for multi-class classification obtained by applying the ‘colorectal-tiatoolbox-resnet50.kather100k’ model on a colon biopsy. Each tile is assigned to the class with the highest prediction score and colored accordingly. (C) Density map obtained by applying the ‘gmblgg-survival-porpoise.tcga’ model on a glioblastoma resection. Here, brighter areas correspond to tiles associated with higher percentile-ranked attention scores.

The integration workflow supports both a default and an on-demand mode (**Figure 3**). In both cases the interaction between the AP-LIS and the AI-DSS initiates in conjunction with a laboratory order message (OML^O33) request generated from the AP-LIS and transmitted to the AI-DSS. In the current implementation, two types of events can trigger the OML^O33 message: (i) the scanning of a new routine slide (default mode), and (ii) a request placed by pathologists through the virtual tray of the AP-LIS (on-demand mode). In the default mode, when a new histopathological glass slide is digitized, and its tissue type and staining match at least one available DL model, the AP-LIS will automatically send analysis requests by triggering as many laboratory order messages as the number of DL models suitable for the tissue type. The default mode runs continuously in the background and does not require any user manual intervention. This operational modality is reserved for all classification models except for those predicting clinical biomarkers status and risk of cancer death. Following an OML^O33 HL7 message, the analysis status can be monitored through the virtual tray of the AP-LIS (**Figure 4A**), where pathologists can visualize whether a slide has already been analyzed or an analysis is in progress. If multiple DL algorithms have been run on the same WSI, opening the slide in the virtual tray will display a pop-up window listing all deployed model names, and pathologists can choose which model results to visualize in QuPath through a double click (**Figure 4B**). Finally, in cases where the analysis fails, as for example when the server-client architecture is not in listening mode and no connection can be established between the AP-LIS and the AI-DSS, the WSI gets labeled with the flag *Analisi fallita* [Analysis failed]. In the on-demand mode the AP-

**Figure 3.**
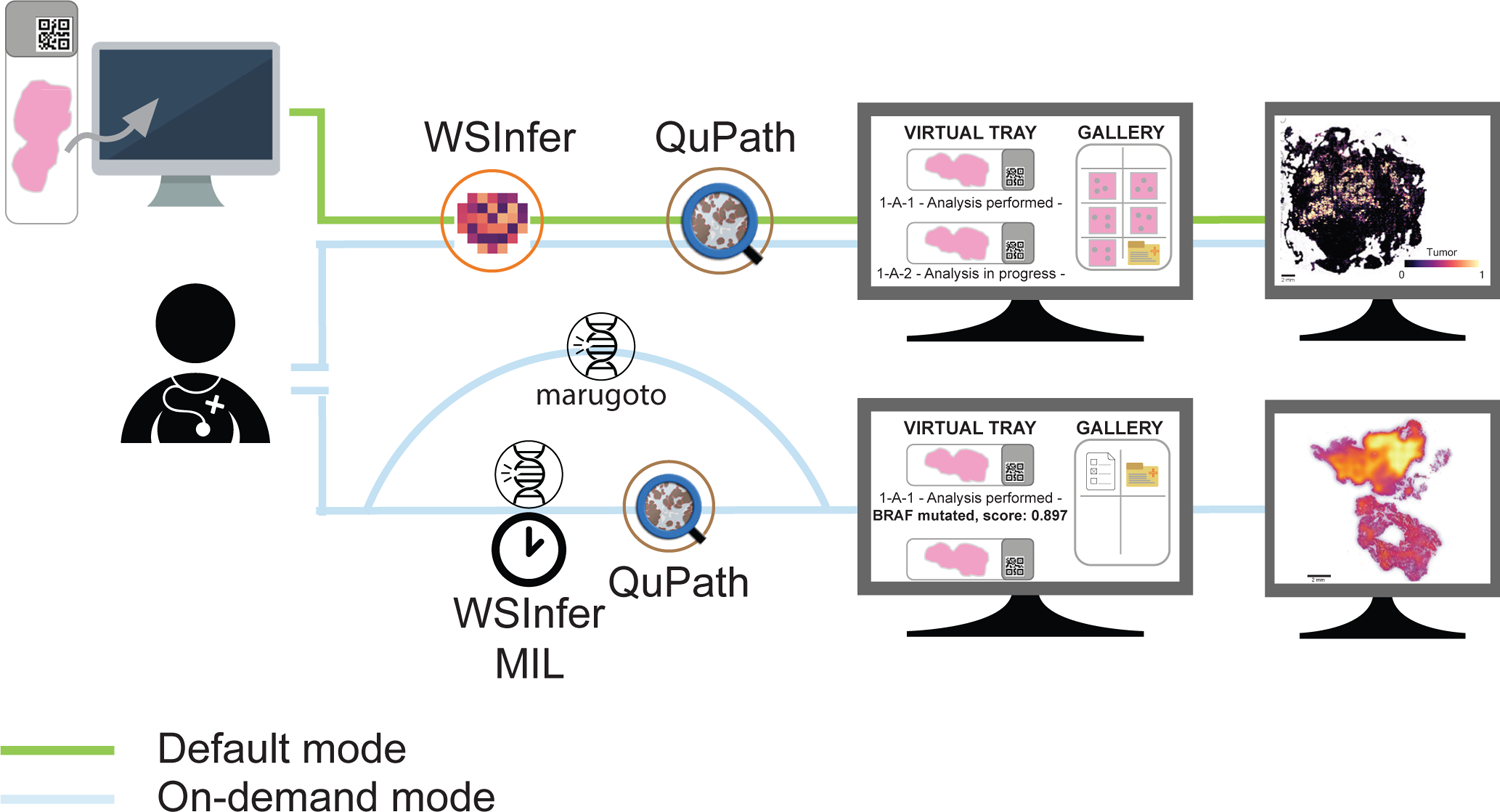
Operational modalities of the integration workflow. In the default mode (green path) new digitized slides are automatically analyzed relying on DL models (both built-in WSInfer and custom developed models) for the classification of different tissue types or conditions. In this modality, WSInfer is used for DL model deployment and QuPath to build colored heatmaps for the visualization of model inference results. For selected models the gallery is populated with the top five predicted tiles supporting the worst clinical outcome. In the on-demand mode (blue path), the analysis request is manually initiated by pathologists from the virtual slides tray. In addition to the models applicable in the default mode, this modality allows applying DL models for prediction of clinical biomarkers and risk of cancer-related death through the WSInfer-MIL and the marugoto toolboxes. Slide-level predictions are displayed as WSI description in the virtual tray. For WSInfer-MIL, visualization of model predictions through density maps are also provided.

**Figure 4.**
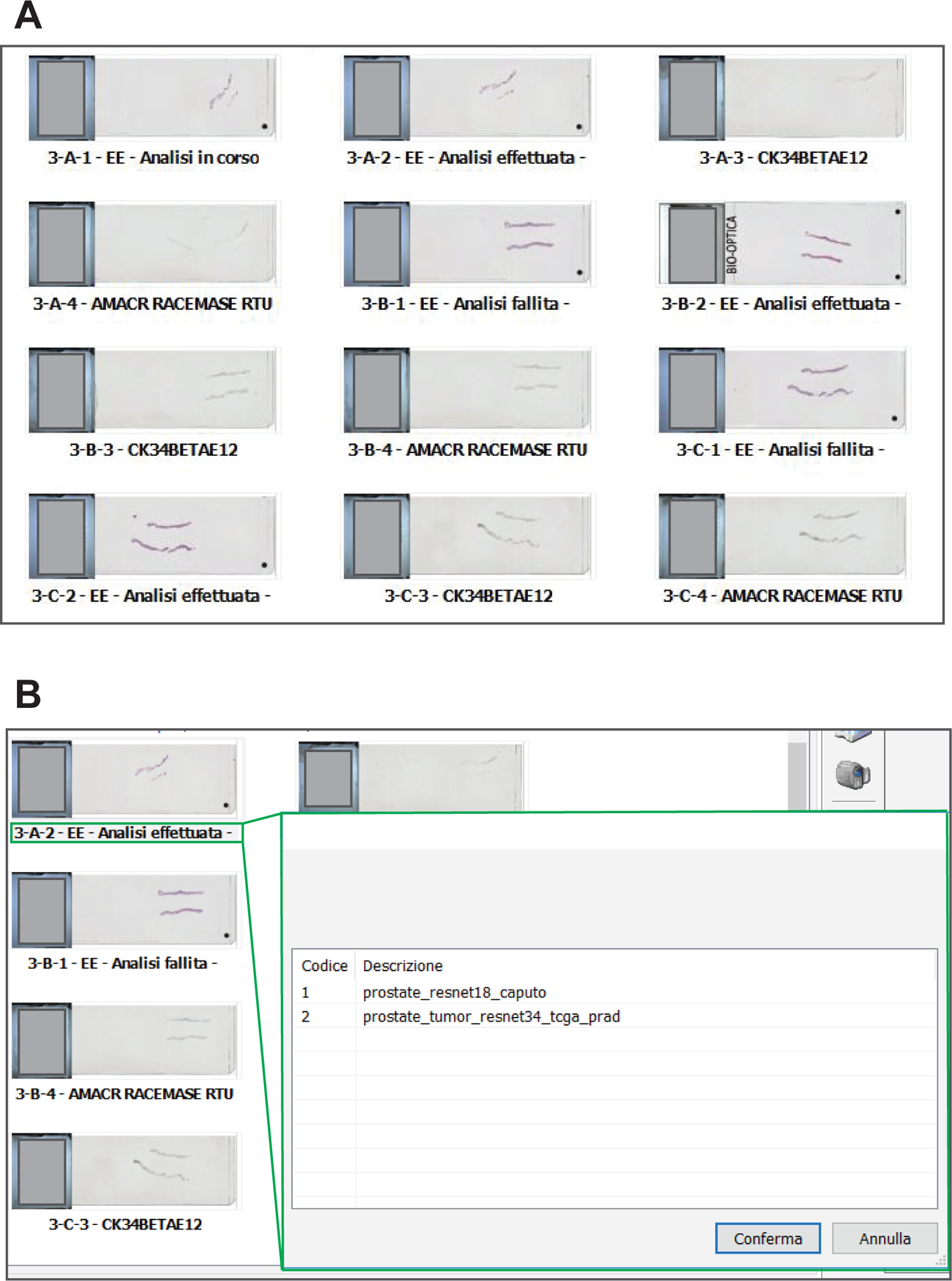
Snapshot of the virtual tray of the AP-LIS following a laboratory order message request. (A) Overview of the virtual tray of the AP-LIS with multiple whole-slide images (WSIs) for the same patient. WSIs for which an analysis request has been sent from the AP-LIS to the DSS, and are currently in a queue waiting for deep-learning (DL) model deployment, are flagged as “Analisi in corso” [Analysis in progress] (i.e., 3-A-1). WSIs already successfully analyzed are flagged as “Analisi effettuata” [Analysis performed] (e.g., 3-A-2). WSIs whose analysis resulted in an error are flagged as “Analisi fallita” [Analysis failed] (e.g., 3-B-1). (B) Pop-up window listing all DL models run on a WSI and selectable for visualization. Annulla [Cancel]; Codice [Code]; Conferma [Confirm]; Descrizione [Description]; EE = Ematossilina – Eosina [Hematoxylin&Eosin].

LIS initiates an order message request for a given WSI only if explicitly triggered by a pathologist from the virtual tray. Pathologists can initiate an analysis request by first right clicking on the WSI to analyze, and then selecting the DL model of choice among the list of available DL models by left double clicking. The on-demand mode is particularly useful to: (i) re-analyze cases assigned at the acceptance stage to a wrong tissue type, and therefore analyzed by default with a wrong DL model; (ii) re-analyze WSIs whose analysis failed in default-mode, because of failing to establish a connection between the AP-LIS and the AI-DSS; (iii) run DL models for the prediction of clinical biomarkers or cancer-related risk death after a confirmed diagnosis of cancer. Taking into account these use-case scenarios, the whole set of DL models integrated in the workflow are available to pathologists for on-demand requests, including those aimed at predicting clinical biomarkers status and risk of cancer death. Analogously to the default mode, also for the on-demand mode the progress status of WSIs analysis is highlighted in the AP-LIS virtual tray.

Once the analysis related to a WSI has been successfully completed, all the clinical-relevant information supporting DL model final predictions are transmitted from the AI-DSS to the AP-LIS as OBX segments via OUL^R21 HL7 messages. Notably, for selected patch-based classification models, the top five predicted tiles supporting the worst clinical outcome are included in the gallery as jpg files (**Figure 5A**). Instead, for slide-level classification models the predicted class (or risk class) and the associated prediction score (or risk score) are imported in the virtual tray and displayed as WSI description (**Figure 5B**).

**Figure 5.**
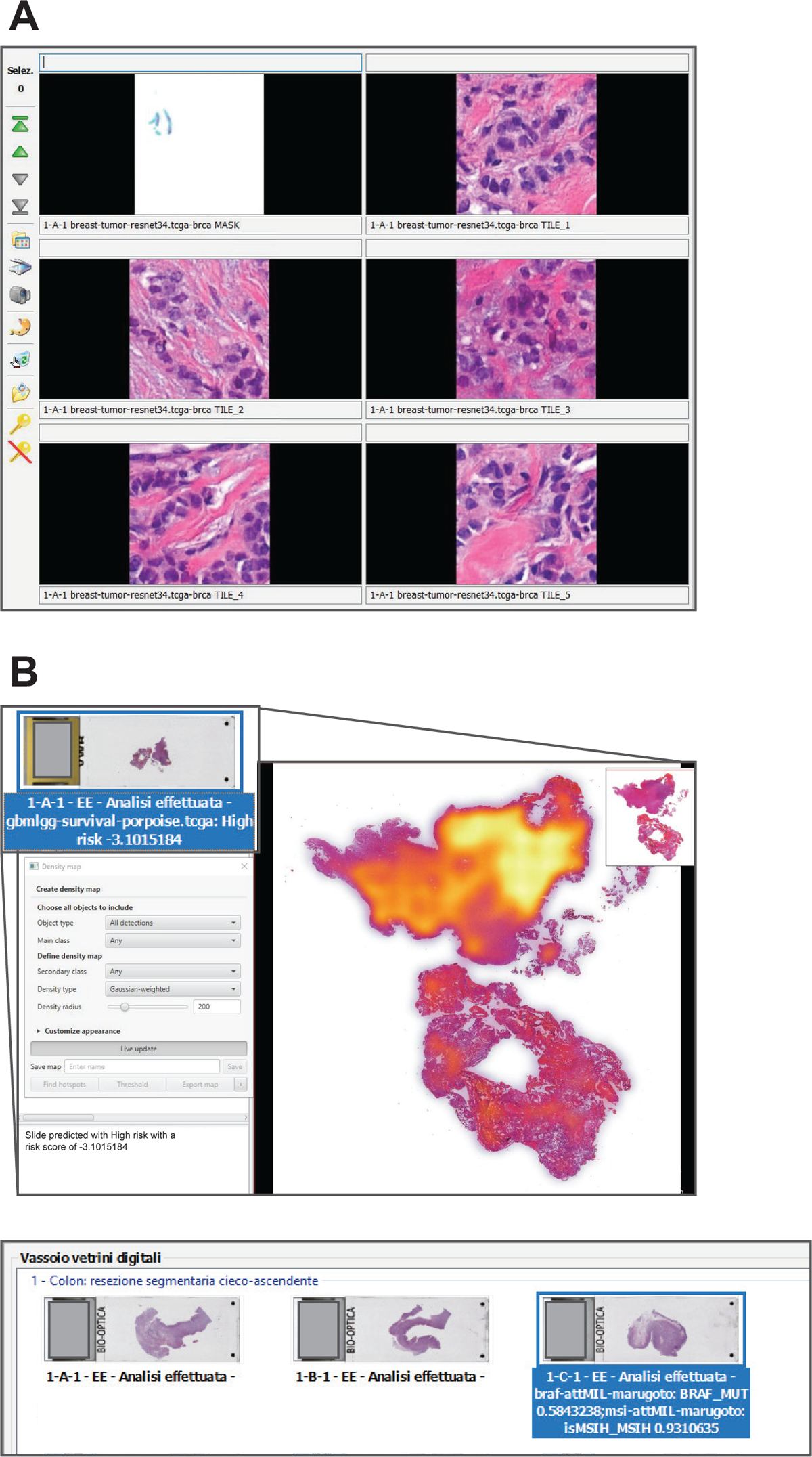
Integration of deep-learning model predictions in the virtual tray. (A) Overview of gallery in the AP-LIS storing the top five predicted tiles supporting the diagnosis of “Tumor” as well as the tissue mask used for patches generation. (B) Visualization of slide-level classification models results in the AP-LIS. (top) WSI analyzed by the ‘gmblgg-survival-porpoise.tcga’ WSInfer-MIL model and predicted with high risk of cancer-related death. The associated density map highlights in yellow tiles associated with higher percentile-ranked attention scores. (bottom) WSI analyzed with both ‘braf-attMIL-marugoto’ and ‘braf-attMIL-marugoto’ models using

In summary, our framework is robust and compatible with several digital pathology tools and data sources, thus potentially applicable to wide range of clinical and research settings.

## Discussion

The potential of AI tools in pathology to enhance diagnosis, prognosis and prediction of clinical biomarkers has been widely acknowledged, marking the onset of a new era often referred to as the ‘third revolution’ in pathology.^23^ However, one of the key factors contributing to the scarce adoption of AI-based solutions in routine settings relies in the difficulty of achieving a full integration of these tools into the anatomic pathology laboratory information system (AP-LIS)^9^.

Here, we leveraged a fully digitized pathology infrastructure^15^ to develop a standardized, open-source Python-based framework that allows the integration of both publicly available and custom developed DL algorithms into the AP-LIS, and the visualization and interpretation of DL model results through colored heatmaps. Importantly, the integration was based on HL7, one of most relevant EHR standards for the exchange of electronic healthcare data between medical information systems^24^.

Furthermore, the use of freely-available resources for both deployment of DL models and visualization of DL model results reduces implementation costs, allows customizing and extending functionalities according to specific needs, and enhances workflow’s portability across different healthcare environments.

WSInfer^13^ and WSInfer-MIL were chosen as main toolboxes for the deployment of patch-level and slide-level pre-trained DL models respectively, since they implement a streamlined, fast, end-to-end pipeline that goes from tissue detection to DL model deployment without requiring any user intervention. The workflow proved able to successfully support different types of classification tasks, from strongly to weakly supervised DL algorithms. Furthermore, the integration of marugoto as additional toolbox for the prediction of *BRAF* mutational status and microsatellite instability (MSI), further highlighted the flexibility and adaptability of the framework towards publicly available resources, in addition to enabling extension of the pool of integrated algorithms for the prediction of predictive clinical biomarkers. The open-source bio-image analysis tool QuPath^19^ was used to visualize DL model results. QuPath had been already fully integrated within the AP-LIS of the Caltagirone digital lab in April 2023 since it not only allows WSIs visualization, but it also supports high-throughput, reliable image analyses, enabling objective quantification of clinically relevant biomarkers (e.g., estrogen receptor, progesterone receptor) in different cancer entities.^25^ One of the major limitations of commercial AI solutions is often the requirement for specific visualization software that does not easily integrate with existing AP-LIS or Picture Archiving and Communication Systems (PACS)^9^, thus ultimately contributing to the slow uptake of AI technologies in the clinical practice. In contrast, QuPath provides a comprehensive visualization capability without the need for multiple software, irrespective of the type of DL algorithm employed.

The integration solution we implemented includes two operational modalities: (i) a default mode where the model is automatically configured to run as each new slide is scanned; and (ii) an on-demand mode where pathologists can initiate the analysis by selecting the model directly from the virtual tray of the AP-LIS. This was exactly what wished for by van Dienst et al., who in their recent review highlighted how the efficiency of AI tools will be fully exploited only when the integrated AI solution can run automatically on the backend or be specifically triggered at need during diagnostics.^9^ In Hartman et al. these two modalities were referred to as ‘preprocessing fashion’ and ‘on-demand fashion’ respectively.^26^ The developed workflow allows the application of multiple algorithms per slide, as well as integrating the output from various algorithms with each other. For instance, analysis for the prediction of *BRAF* mutational status can be requested on-demand by pathologists after a confirmed diagnosis of cancer based on a previously applied classification model.

Collectively, our study demonstrates that implementing CPath solutions into a fully digitized pathology department is feasible and can be optimized to be highly user-friendly for pathologists. Our framework though, despite its strengths and novelty, faces limitations. Firstly, it is worth emphasizing that all software, except for the AP-LIS, and DL models employed in the study are non-commercial, open-source resources intended for research use only. Hence, use of the integration workflow, with the included software and DL algorithms, outside of research context is under the responsibility of the user. Indeed, in order for a CPath solution to be deployed for clinical use, rigorous validation and regulatory approval (e.g., FDA or IVD) are crucial requirements. These requirements are even more important when considering ethical issues^27^ raising from the application of AI tools to WSIs as well as possible legal liability consequences^28,29^ if errors occur.

In addition, digitization of the pathology workflow is an absolute requirement for the application of our integration framework. Interoperability with diverse digital pathology systems and varying computing infrastructures requires an initial setup with potential customizations that can impact implementation efficiency. Furthermore, pathologists’ acceptance and proficiency in utilizing AI tools represent other barriers, necessitating ongoing training and support initiatives. Possible pitfalls and caveats are numerous and include biases inherent in the model (e.g., lack of data diversity, poor generalizability) but also downstream issues such as pathologist deskilling.^30^

In conclusion, the described workflow provides a proof-of-concept framework for a standardized, portable, and open-source solution able to run DL algorithms routinely within the AP-LIS relying on HL7 messaging and freely available software for digital pathology. This solution allows, for the first time, to effectively close the gap between the research area and the clinical implementation of AI-based tools, thus marking a first step towards their ‘realistic assessment’^31^ in routine settings. Continued innovation in DL algorithms and AI model interpretability will reinforce our commitment to advancing AI-driven diagnostics in pathology, ultimately improving patient outcomes and healthcare delivery.

## Contributors

MA, FuFe, and FiFr conceived the study, interpreted results, and wrote the manuscript. MA implemented the integration framework. DR embedded results in Pathox. SS, AC, FM, and AH contributed materials, clinical, and/or methodological expertise. FuFe and FiFr supervised the work. All authors edited the final manuscript. All authors accept the final responsibility to submit for publication and take responsibility for the contents of the manuscript.

## Declaration of interests

DR is full employee of TESI Group/GPI. AH receives honoraria for lectures or consulting/advisory boards for Abbvie, Agilent, AstraZeneca, Biocartis, BMS, Boehringer Ingelheim, Cepheid, Diaceutics, Gilead, Illumina, Ipsen, Janssen, Lilly, Merck, MSD, Nanostring, Novartis, Pfizer, Qiagen, QUIP GmbH, Roche, Sanofi, 3DHistotech and other research support from AstraZeneca, Biocartis, Cepheid, Gilead, Illumina, Janssen, Nanostring, Novartis, Owkin, Qiagen, QUIP GmbH, Roche, Sanofi. AC, FM and FiFr report ad hoc advisory board membership with Roche Diagnostics, Italia, unrelated to the current work. FiFr is one of the inventors of ‘Sample imaging and imagery archiving for imagery comparison Merlo, P.T. et al. US patent 16/688/613 2020’. The remaining authors have no conflicts of interest to declare.

## Acknowledgments

Deutsche Forschungsgemeinschaft (German Research Foundation) grant TRR 305 [project Z01 (FuFe)]; European Union-Next Generation EU-NRRP M6C2-Investment 2.1 Enhancement and strengthening of biomedical research in the NHS (DIPLOMAT-PNRR-MR1-2022-12375735) [FiFr]; Bavarian Funding Programme for the Initiation of International Projects (BayIntAn) [FuFe and FiFr].

## Pre-trained DL models included in the integration workflow

For each DL model a reference to the relevant publication(s) and/or model’s repository (column ‘Reference’), a summary description of the task performed (column ‘Model Description’), the classes provided in output (column ‘Classes’), the type of visualization heatmap chosen (column ‘Visualization’), and the toolbox used for model deployment (column ‘Toolbox’) are provided. GMBLGG = Low-Grade Glioma; HPV = Human Papillomavirus; KIRP = Kidney Renal Papillary Cell Carcinoma; MET = Metastasis; MSI = Microsatellite instability; MUT = Mutated; NOMET = Non Metastasis; TIL = Tumor Infiltrating Lymphocytes; WT = Wild Type.

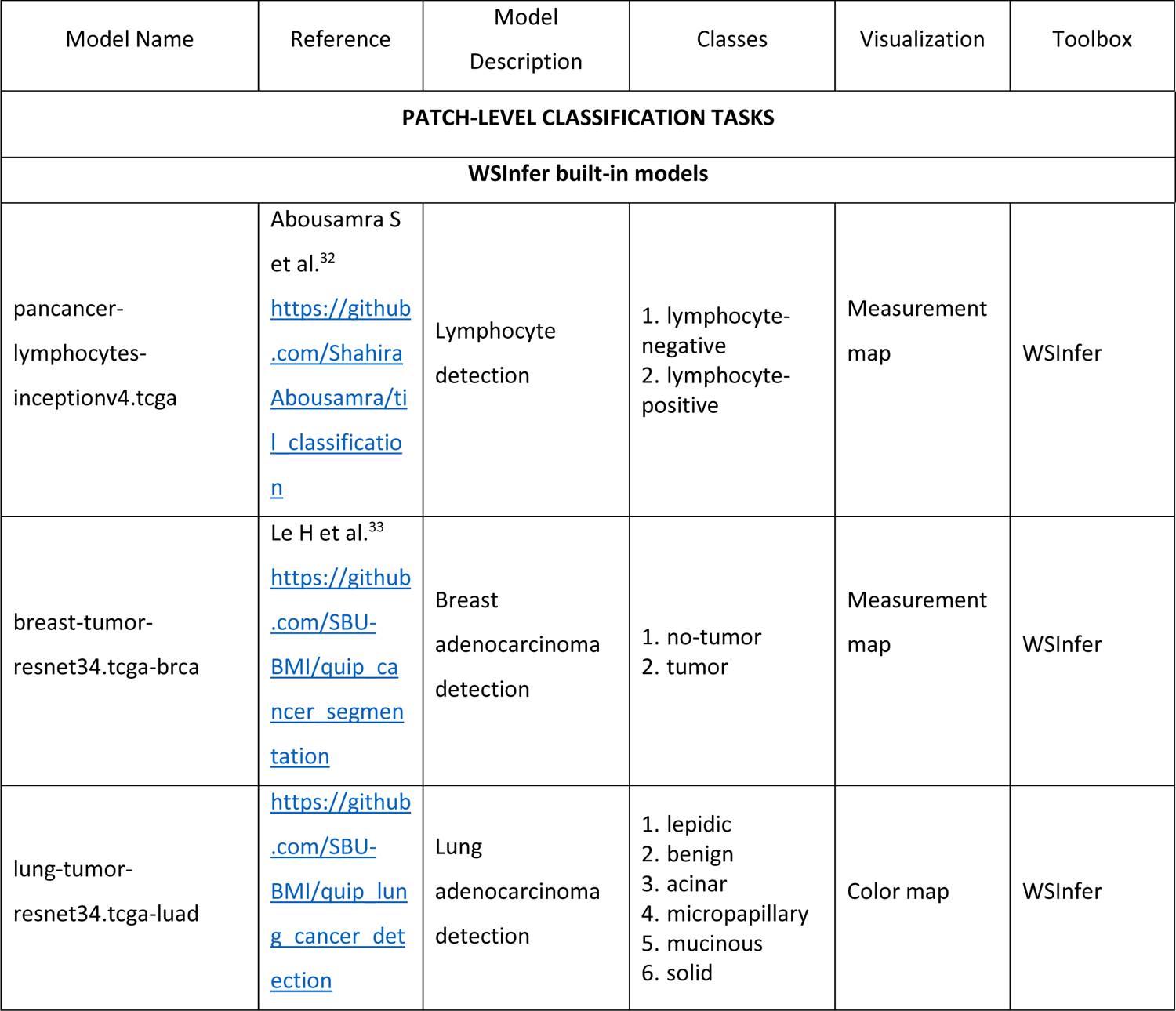
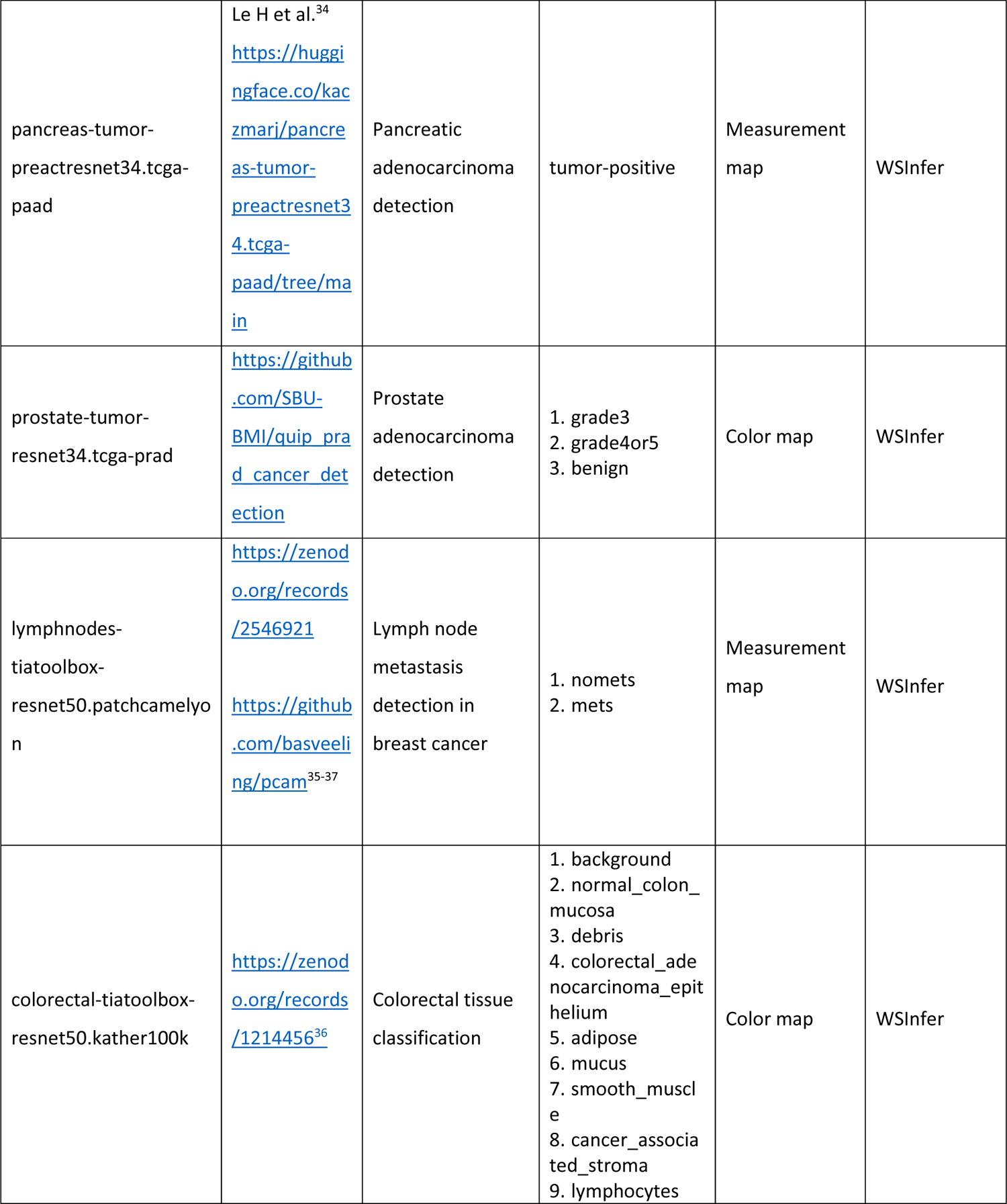
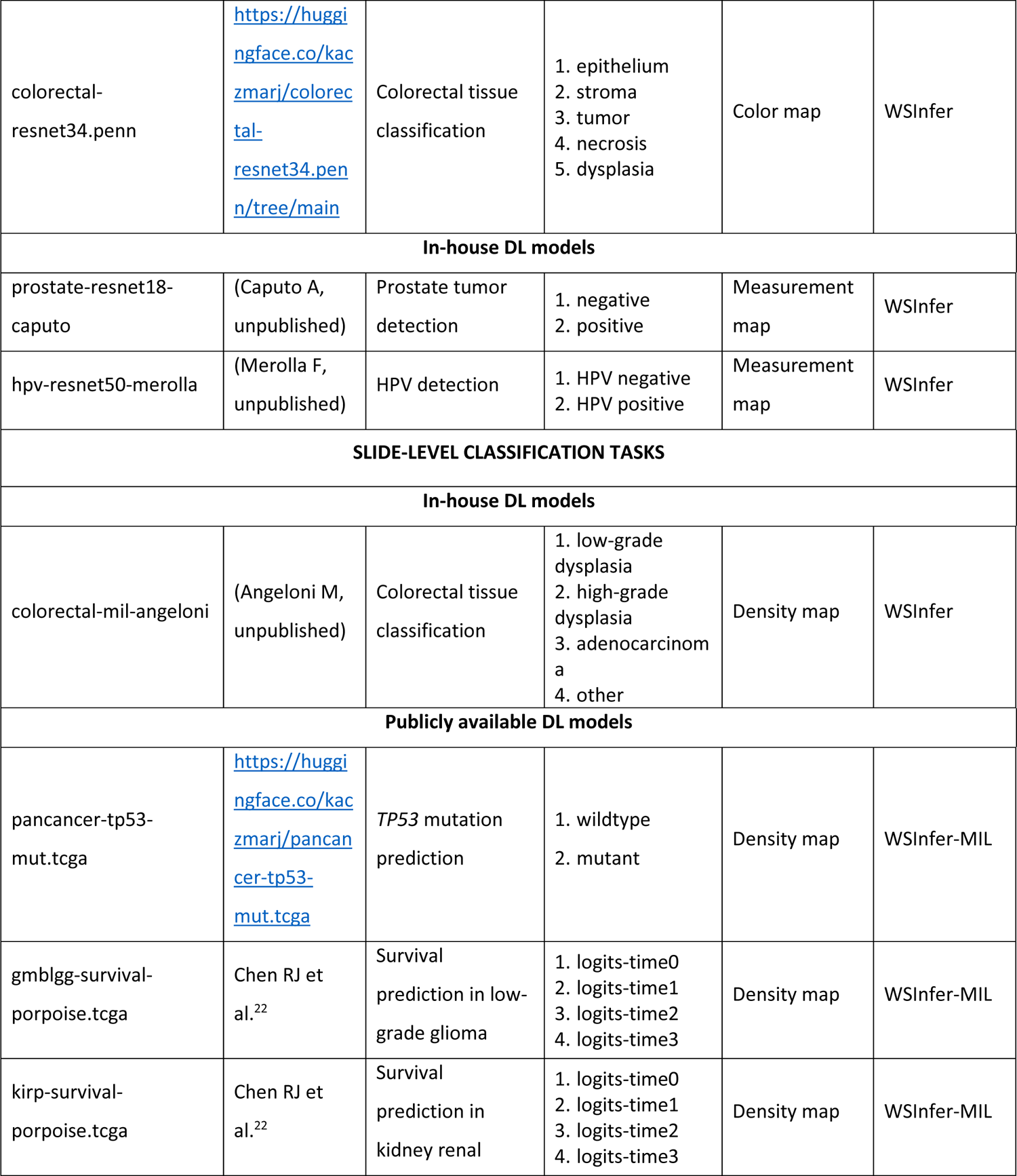
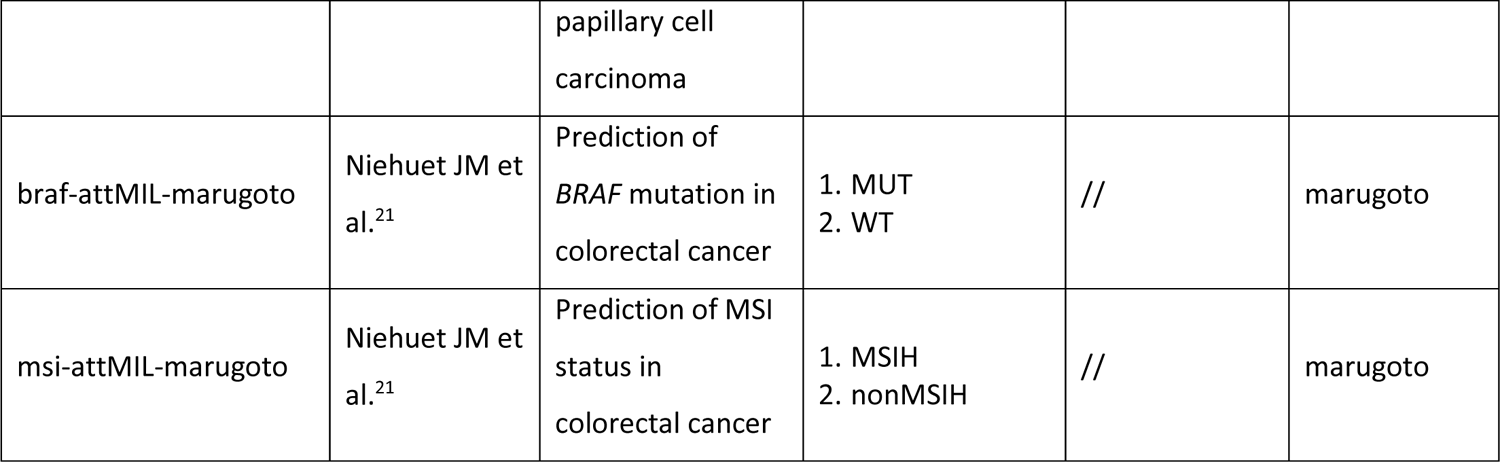

**Supplementary Figure 1.**
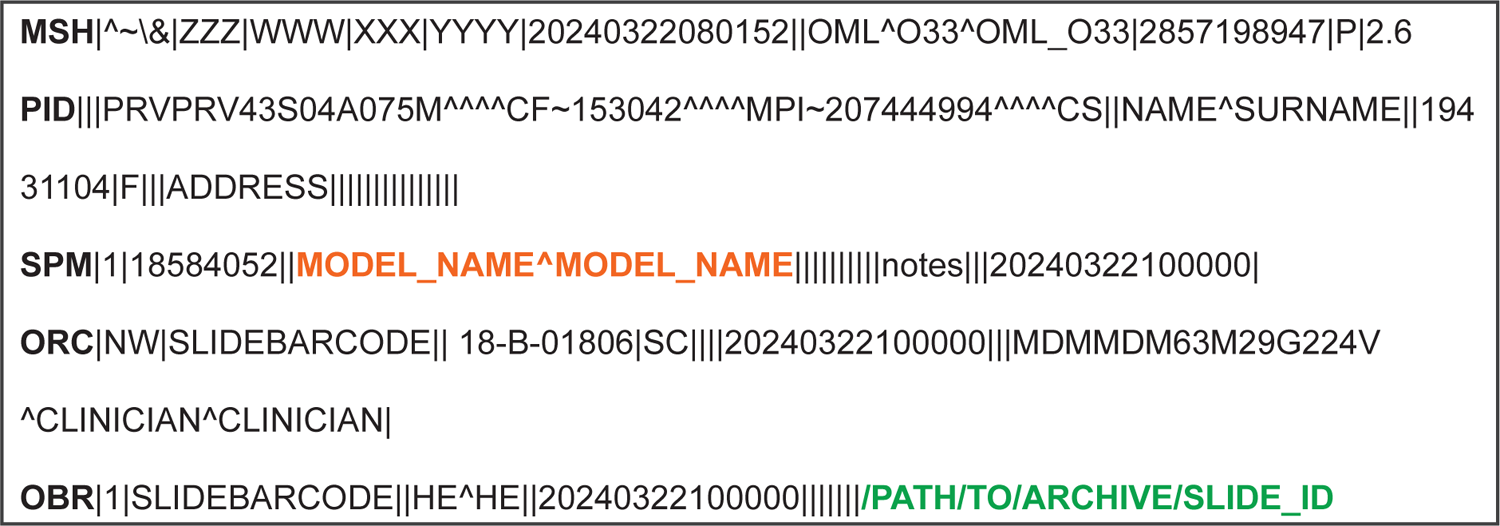
Example of OML^O33 HL7 message. Sample laboratory order HL7 version 2.6 message transmitted from the AP-LIS to the AI-DSS for the analysis of a WSI. Each OML^O33 message is made up of five segments, i.e., MSH, PID, SPM, ORC, and OBR (bold black). The name of the DL model to deploy is stored in the fields 4.1 and 4.2 of the SPM segment (bold orange), whereas the path to the WSI to analyze is retrieved from field 13 of the OBR segment (bold green).

**Supplementary Figure 2.**
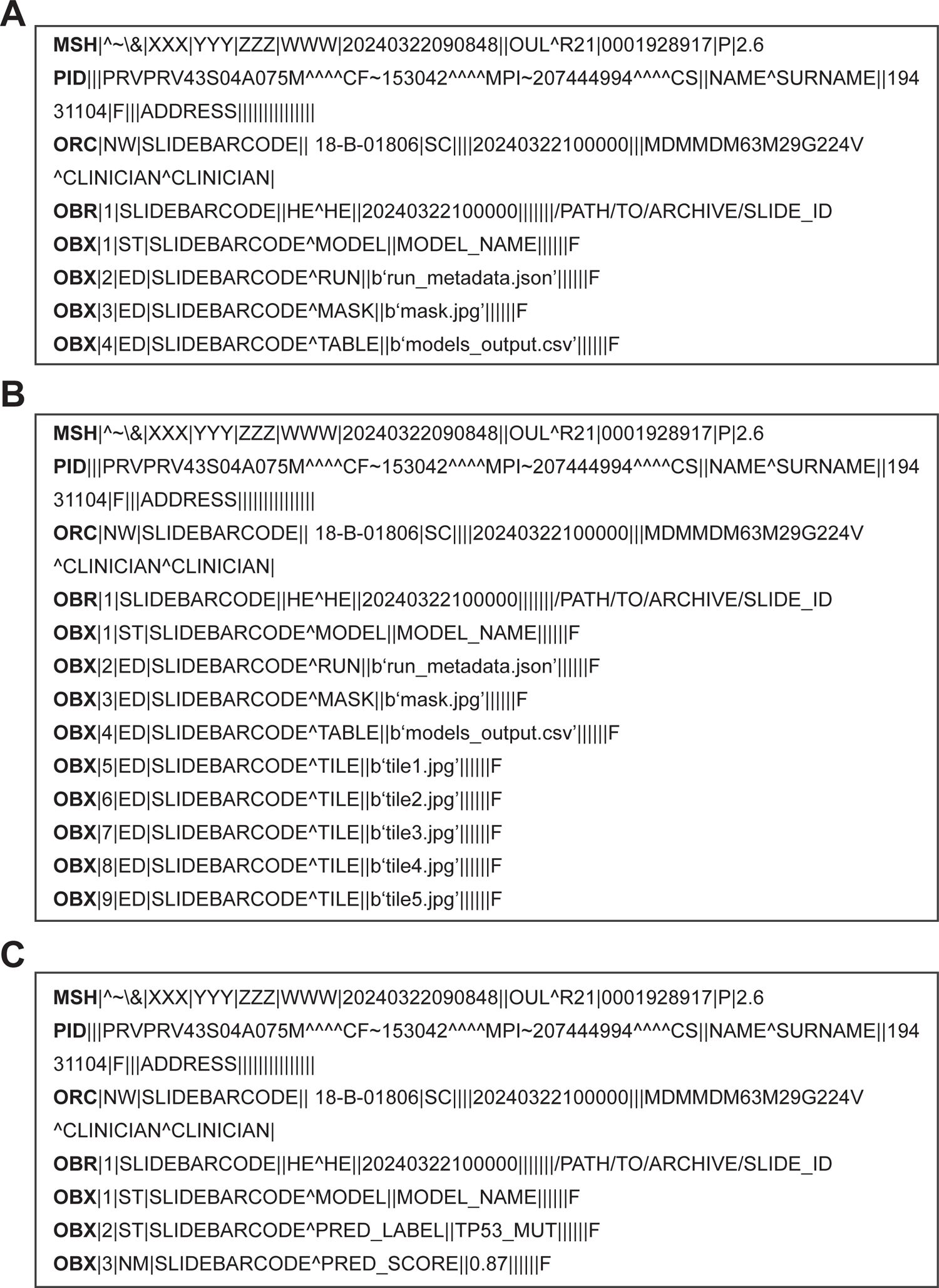
Examples of OUL^R21 HL7 messages. Sample unsolicited laboratory observation HL7 version 2.6 messages transmitted from the AI-DSS to the AP-LIS after (A) running model deployment with the WSInfer CLI tool, (B) running model deployment with the WSInfer CLI tool for a subset of classification tasks, and (C) running model deployment with pre-trained slide-level classification models.

## Videos legend

**Video 1. Activation of measurement map in QuPath.** Measurement map visualization is enabled by (1) navigating to Measure → Show measurement map (or alternatively pressing simultaneously Ctrl+Shift+M), (2) selecting the label measurement (e.g. Tumor) popping up in the Measurement map window, (3) toggling on the detection objects through the ‘Show/hide detection objects’ toolbar button, and (4) filling up the detection objects through the ‘Fill/Unfill detection object ROIs for display’ toolbar button.

**Video 2. Activation of color map in QuPath.** Color map visualization is enabled by filling up the detection objects through the ‘Fill/Unfill detection object ROIs for display’ toolbar button and the displayed classes can be visualized navigating to the ‘Annotations’ tab.

**Video 3. Activation of density map in QuPath.** Density map visualization is enabled by (1) navigating to Analyze ◊ Density maps ◊ Create density map. (2) Density map appearance can be modified by changing the density type to, e.g., Gaussian-weighted. Slide-level outcome is saved as free text in the white box on the bottom left panel under the Project tab.

## Supplementary Materials and Methods

### Deployment of patch-level and slide-level pre-trained deep-learning models through WSInfer and WSInfer-MIL

Deployment of pre-trained patch-level classification models was performed relying on the WSInfer^13^ command-line (CLI) tool v0.6.1. WSInfer was installed in the dedicated conda environment via pip after PyTorch installation, as recommended by the authors (https://github.com/SBU-BMI/wsinfer; https://wsinfer.readthedocs.io/en/latest/installing.html). The tool provides a fast, end-to-end workflow that: (i) segments the tissue in the WSI, (ii) creates patches of the segmented tissue region, (iii) runs the pre-trained model on the extracted patches, and (iv) saves model results, with minimal, if any, user intervention. WSInfer allows the deployment of both built-in and customized DL models. Pre-trained weights for built-in models, with the associated configuration files, were retrieved relying on the Python package wsinfer-zoo v0.6.2 (https://github.com/SBU-BMI/wsinfer-zoo; https://zenodo.org/records/12680690), automatically installed as dependency during the WSInfer pip installation. Deployment of pre-trained DL models on routine WSIs was performed by running the wsinfer run command with openslide v3.4.1 as backend for loading WSIs (--backend=openslide argument) and relying on the default settings for both batch size (--batch-size argument) and number of workers (--num-workers argument). For WSInfer built-in models, the wsinfer run command was launched using the following required arguments: a directory of WSIs (--wsi-dir argument), a directory where to store results (--results-dir argument), and the name of the DL model to deploy (--model argument). For inference with models not included in WSInfer, the model to deploy was instead specified by adding to the wsinfer run command the path to the local model configuration json file (--config argument), and the path to the pre-trained local model in TorchScript format (--model-path argument). The wsinfer run command creates, by default, a structured output folder storing, for each WSI: a tissue mask, an h5 file containing patch coordinates, a json file storing the metadata of the run, and a CSV file storing model inference results. The latter contains a row for each tile, with columns storing the x and y coordinates of the tiles in base pixel units, their height and width, and their prediction score, one for each class. All the built-in models provided by the WSInfer CLI tool were included in the integration workflow.

Deployment of pre-trained attention-based Multiple Instance Learning (MIL) models for slide-level classification tasks was performed relying on the WSInfer-MIL CLI tool v0.1.0 (https://github.com/SBU-BMI/wsinfer-mil; https://zenodo.org/records/12680704), which was installed on the dedicated conda environment via pip following authors’ instructions. Analogously to the WSInfer patch-level counterpart, wsinfer-mil allows deploying both built-in and customized DL models. Inference of built-in models was performed running the wsinfer-mil run command with the following required arguments: the HuggingFace Hub repository ID of the model (--hf-repo-id argument), and the path to the WSI (--wsi-path argument). To run model inference on mrxs files, the modules wsi_utils.py, inference.py, and data.py of the wsinfer-mil package were modified to replace tiffslide with openslide for WSIs opening and reading. Furthermore, differently from the WSInfer CLI-tool, WSInfer-MIL outputs model predictions directly to the command line interface, without saving them in any external file format. To streamline the post-processing of DL model inference results, the output_container.py and the inference.py modules were further modified to save both slide-level and tile-level predictions in CSV format. In the CSV file with slide-level predictions, the number of rows was taken equal to the number of classes, with the first column storing the class names, and the second column containing the prediction probability for each class. In the CSV file with tile-level predictions, the number of rows was instead taken equal to the number of patches, and six columns were created to store for each patch the x coordinate, the y coordinate, the width, the height, the attention score, and the percentile-ranked attention score. Percentile ranked attention scores were calculated, starting from the un-normalized attention scores, using the Pandas rank() method and setting to true the argument ‘pct’. The following WSInfer-MIL built-in models were included in the integration workflow: pancancer-tp53-mut.tcga (https://huggingface.co/kaczmarj/pancancer-tp53-mut.tcga), gbmlgg-survival-porpoise.tcga (https://huggingface.co/kaczmarj/gbmlgg-survival-porpoise.tcga), and kirp-survival-porpoise.tcga (https://huggingface.co/kaczmarj/kirp-survival-porpoise.tcga). For survival models, the formula to calculate the risk score basing on model output logits, as well as the median risk score to split patients into low risk and high risk, were retrieved from the respective model’s huggingface.com page.

### Deployment framework for the assessment of BRAF mutational status and MSI

Pre-trained models for the assessment of BRAF mutational status and microsatellite instability (MSI) were retrieved from the GitHub repository made public by Niehues et al.^21^ (https://github.com/KatherLab/crc-models-2022/tree/main/Quasar_models). For both classification tasks, the export-0.pkl file under the corresponding Wang+attMIL subfolder was chosen for model inference on new WSIs.

The two models were deployed using the marugoto toolbox v.0.8.0 (https://github.com/KatherLab/marugoto). marugoto was installed in the same conda environment hosting WSInfer and WSInfer-MIL CLI tools by first cloning the GitHub repository https://github.com/KatherLab/marugoto and then running the command ‘pip install.’ from the marugoto subfolder, as specified in the documentation (https://github.com/KatherLab/marugoto/blob/main/Documentation.md). Similar to WSInfer-MIL, marugoto allows the construction of DL workflows for weakly supervised classification problems: from features extraction to model deployment on pre-extracted features. However, differently from the WSInfer and WSInfer-MIL CLI tools, tissue segmentation and patches generation are not included in the workflow. Instead, the workflow assumes that features extraction is applied to previously extracted patches saved as jpeg files. To make tissue segmentation and patches generation as efficient as possible, and to avoid saving patches as jpeg files, the module xiyue_wang for feature extraction was customized to take in input the h5 file containing the coordinates of the patches generated by the WSInfer CLI tool. Notably, all WSIs analyzed with marugoto underwent first tissue segmentation and patches generation through the WSInfer command *wsinfer patch*, and then the output h5 file with patches coordinates was used as input for running the marugoto.extract.xiyue_wang features extraction module. To generate patches coordinates, the wsinfer patch command was run using the following required arguments: a directory of WSIs (--wsi-dir argument), a directory where to store results (--results-dir argument), the patch size in pixel (--patch-size-px argument), and the physical spacing of the patches in mpp (--patch-spacing-um-px argument). A patch size of 224 pixel and a patch spacing value of 1.14 mpp were used for patches generation.^21^ Finally, model deployment on the pre-extracted features was performed relying on the marugoto.mil deploy module. The module was run using the target label ‘BRAF’ (--target_label argument) and the categorical labels [’MUT’,’WT’] (--cat_labels argument) for the assessment of BRAF mutational status, and using the target label ‘isMSIH’ and the categorical labels [’MSIH’,’nonMSIH’] for the assessment of MSI. The clinical table (--clini_table argument) was created to contain one column called ‘PATIENT’, storing patient identifier, and one column named as the target label (i.e., BRAF or isMSIH) containing either of the two associated categorical labels. The slide table csv file (--slide-csv argument) was instead created to contain one column called ‘PATIENT’, storing the same patient identifier found in the clinical table, and one column called “FILENAME” with the filename, without extension, of the feature vectors h5 file.

### Visualization of deep-learning model inference results in QuPath

The generation of tiled colored overlays as visual representation of DL model inference results was performed relying on the open-source bio-image analysis software QuPath^19^ v0.4.3 and on the Python package paquo v0.8.0 (https://github.com/Bayer-Group/paquo – copyright 2020 Bayer AG, licensed under GPL-3.0) that provides a pythonic interface to create and work with QuPath projects.

For each analyzed WSI, a new QuPath project (.qpproj file) was instantiated using the QuPathProject class from the paquo.projects module and populated with the WSI through the QuPathProject.add_image() method. Tile detections were added to the project hierarchy using the QuPathPathObjectHierarchy.add_tile() method from the paquo.hierarchy module. Each tile was drawn as a polygon using the Polygon.from_bounds() method of the Python package shapely v2.0.4. To correctly build-up tile detection objects in QuPath, the openslide patch coordinates generated by WSInfer and WSInfer-MIL CLI tools needed to be offset. To this aim, an x and y offset were automatically extracted for each WSI through the openslide.bounds-x and openslide.bounds-y properties, and subtracted from the original patch coordinates. The offset coordinates were then used as input for the Polygon.from_bounds() class method. Three different visualization styles were implemented, namely: measurement maps, color maps, and density maps.

In measurement maps, each tile was added to the project hierarchy using the polygon defined with Polygon.from_bounds() as value for the parameter ‘roi’, the worst label from the clinical point of view among DL model categories as value for the parameter ‘path_class’, and the corresponding prediction score as value for the parameter ‘measurements’.

In color maps, a number of new classes equal to the number of categories of the DL algorithm, each associated with a color pair, were defined within the QuPath project. The new classes were instantiated using the QuPathPathClass class from the paquo.projects module and assigned to the project through the QuPathProject.path_class property. Tiles were added to the project hierarchy analogously to measurement maps, this time using as values for the ‘path_class’ and ‘measurements’ parameters the corresponding predicted label, and the associated prediction score, determined by taking the argmax of the predictions across all classes.

For density maps, in addition to tile detection objects, we created in correspondence of each tile coordinates a number of detection points proportional to its percentile-ranked attention score. Notably, nine detections were drawn for scores >= 0.9, eight detections for scores between 0.9 and 0.8, and so forth until reaching a minimum of one detection point for percentile-ranked attention scores lower than 0.2. Detection points were created relying on the Point and MultiPoint functions of the shapely package and added to the project hierarchy through the QuPathPathObjectHierarchy.add_detection() method from the paquo.hierarchy module.

Furthermore, a brief description of DL model inference results including the slide-level predicted label/risk class and the associated predicted score/risk score, was added to the QuPah project as free text through the QuPathProjectImageEntry.description property of the paquo.hierarchy module.

## References

1. Kiran N, Sapna F, Kiran Fet al. Digital pathology: transforming diagnosis in the digital age. Cureus 2023; 15.

2. Treanor D. Virtual slides: an introduction. Diagnostic histopathology 2009; 15: 99–103.

3. Geaney A, O’Reilly P, Maxwell P, James JA, McArt D, Salto-Tellez M. Translation of tissue-based artificial intelligence into clinical practice: from discovery to adoption. Oncogene 2023; 42: 3545–55.

4. Unger M, Kather JN. Deep learning in cancer genomics and histopathology. Genome Medicine 2024; 16: 44.

5. Asif A, Rajpoot K, Graham S, Snead D, Minhas F, Rajpoot N. Unleashing the potential of AI for pathology: challenges and recommendations. The Journal of Pathology 2023; 260: 564–77.

6. Markowetz F. All models are wrong and yours are useless: making clinical prediction models impactful for patients. npj Precision Oncology 2024; 8: 54.

7. Reis-Filho JS, Kather JN. Overcoming the challenges to implementation of artificial intelligence in pathology. JNCI: Journal of the National Cancer Institute 2023; 115: 608–12.

8. Verghese G, Lennerz JK, Ruta D et al. Computational pathology in cancer diagnosis, prognosis, and prediction–present day and prospects. The Journal of Pathology 2023; 260: 551–63.

9. van Diest PJ, Flach RN, van Dooijeweert C et al. Pros and cons of artificial intelligence implementation in diagnostic pathology. Histopathology 2024; 84: 924–34.

10. Flach RN, Stathonikos N, Nguyen TQ, Ter Hoeve ND, van Diest PJ, van Dooijeweert C. CONFIDENT-trial protocol: a pragmatic template for clinical implementation of artificial intelligence assistance in pathology. BMJ open 2023; 13: e067437.

11. Wagner SJ, Matek C, Boushehri SS et al. Built to last? Reproducibility and reusability of deep learning algorithms in computational pathology. Modern Pathology 2024; 37: 100350.

12. Wagner SJ, Matek C, Shetab Boushehri S et al. Make deep learning algorithms in computational pathology more reproducible and reusable. Nature Medicine 2022; 28: 1744–6.

13. Kaczmarzyk JR, O’Callaghan A, Inglis F et al. Open and reusable deep learning for pathology with WSInfer and QuPath. NPJ Precision Oncology 2024; 8: 9.

14. Caputo A, Macrì L, Gibilisco F et al. Validation of full-remote reporting for Cervicovaginal Cytology: the Caltagirone-Acireale distributed lab. Journal of the American Society of Cytopathology 2023; 12: 378–85.

15. Fraggetta F, Caputo A, Guglielmino R, Pellegrino MG, Runza G, L’Imperio V. A survival guide for the rapid transition to a fully digital workflow: the “Caltagirone example”. Diagnostics 2021; 11: 1916.

16. Kabachinski J. What is Health Level 7? Biomed Instrum Technol 2006; 40: 375–9.

17. Sepulveda JL, Young DS. The ideal laboratory information system. Archives of Pathology and Laboratory Medicine 2013; 137: 1129–40.

18. Sinard JH, Castellani WJ, Wilkerson ML, Henricks WH. Stand-alone laboratory information systems versus laboratory modules incorporated in the electronic health record. Archives of Pathology and Laboratory Medicine 2015; 139: 311–8.

19. Bankhead P, Loughrey MB, Fernández JA et al. QuPath: Open source software for digital pathology image analysis. Scientific reports 2017; 7: 1–7.

20. Augustine TN. Weakly-supervised deep learning models in computational pathology. Ebiomedicine 2022; 81.

21. Niehues JM, Quirke P, West NP et al. Generalizable biomarker prediction from cancer pathology slides with self-supervised deep learning: A retrospective multi-centric study. Cell reports Medicine 2023; 4.

22. Chen RJ, Lu MY, Williamson DF et al. Pan-cancer integrative histology-genomic analysis via multimodal deep learning. Cancer Cell 2022; 40: 865–78. e6.

23. Salto-Tellez M, Maxwell P, Hamilton PW. Artificial intelligence-the third revolution in pathology. Histopathology 2018.

24. Eichelberg M, Aden T, Riesmeier J, Dogac A, Laleci GB. A survey and analysis of electronic healthcare record standards. Acm Computing Surveys (Csur) 2005; 37: 277–315.

25. Loughrey MB, Bankhead P, Coleman HG et al. Validation of the systematic scoring of immunohistochemically stained tumour tissue microarrays using QuPath digital image analysis. Histopathology 2018; 73: 327–38.

26. Hartman DJ. Applications of Artificial Intelligence in Lung Pathology. Surgical Pathology Clinics 2023.

27. Sorell T, Rajpoot N, Verrill C. Ethical issues in computational pathology. Journal of Medical Ethics 2022; 48: 278–84.

28. Cestonaro C, Delicati A, Marcante B, Caenazzo L, Tozzo P. Defining medical liability when artificial intelligence is applied on diagnostic algorithms: a systematic review. Frontiers in Medicine 2023; 10: 1305756.

29. Jones C, Thornton J, Wyatt JC. Artificial intelligence and clinical decision support: clinicians’ perspectives on trust, trustworthiness, and liability. Medical law review 2023; 31: 501–20.

30. Nakagawa K, Moukheiber L, Celi LA et al, editors. AI in pathology: what could possibly go wrong? Seminars in Diagnostic Pathology; 2023: Elsevier.

31. Van der Laak J, Litjens G, Ciompi F. Deep learning in histopathology: the path to the clinic. Nature medicine 2021; 27: 775–84.

32. Abousamra S, Gupta R, Hou L et al. Deep learning-based mapping of tumor infiltrating lymphocytes in whole slide images of 23 types of cancer. Frontiers in oncology 2022; 11: 806603.

33. Le H, Gupta R, Hou L et al. Utilizing automated breast cancer detection to identify spatial distributions of tumor-infiltrating lymphocytes in invasive breast cancer. The American journal of pathology 2020; 190: 1491–504.

34. Le H, Samaras D, Kurc T, Gupta R, Shroyer K, Saltz J, editors. Pancreatic cancer detection in whole slide images using noisy label annotations. Medical Image Computing and Computer Assisted Intervention–MICCAI 2019: 22nd International Conference, Shenzhen, China, October 13–17, 2019, Proceedings, Part I 22; 2019: Springer.

35. Bejnordi BE, Veta M, Van Diest PJ et al. Diagnostic assessment of deep learning algorithms for detection of lymph node metastases in women with breast cancer. Jama 2017; 318: 2199–210.

36. Pocock J, Graham S, Vu QD et al. TIAToolbox as an end-to-end library for advanced tissue image analytics. Communications medicine 2022; 2: 120.

37. Veeling BS, Linmans J, Winkens J, Cohen T, Welling M, editors. Rotation equivariant CNNs for digital pathology. Medical Image Computing and Computer Assisted Intervention–MICCAI 2018: 21st International Conference, Granada, Spain, September 16-20, 2018, Proceedings, Part II 11; 2018: Springer.

